# Climate-driven decoupling of a coevolved mutualism between Joshua trees (*Yucca brevifolia* and *Y. jaegeriana*, Asparagaceae) and pollinating yucca moths (*Tegeticula synthetica* and *T. antithetica*, Prodoxidae)

**DOI:** 10.64898/2026.10.14.759266

**Authors:** Pryce W. Millikin, Jeremy B. Yoder

## Abstract

Environmental change can disrupt ecologically important mutualisms, if coadapted partners respond to changes differently—but modeling differential responses by specialized mutualists can be challenging. Here, we use TARDIS (Temporal Analysis of Reproduction Distributed in Space) to examine how annual weather variation drives flowering in Joshua trees (*Yucca brevifolia* and *Y. jaegeriana*; Asparagaceae), and subsequent fruit production that relies on obligate yucca moth pollinators (*Tegeticula antithetica* and *T. synthetica*; Prodoxidae). We used more than 16,000 observations of Joshua trees and corresponding weather records, covering an 18-year period, to train Bayesian additive regression tree models of flowering and fruiting activity. The models find flowering and fruiting respond differently to temperature and inter-annual precipitation variation, and a hindcast for the years 1900–2025 reveals that years with conditions favoring flowering have become slightly more frequent, while conditions favoring fruiting—and moth activity—have been stable or declining. Trends in this predicted mismatch between Joshua tree flowering and yucca moth activity vary with latitude and elevation, but do not differ in regions identified as climate refugia. Overall, our results suggest climate change to date has had pervasive, if so far small, impacts on both these specialized mutualists.

## Introduction

Interactions between species—mutualistic, antagonistic, or commensal—play out in variable environments that may alter their outcomes and even the identity of the players (Briscoe Runquist et al., 2020; Heath and Tiffin, 2007; Thompson, 2005; Vale et al., 2011). One element of this context-dependence is the prospect that interacting species may respond to variation in their shared environment in differing ways. Describing interacting species’ independent responses to environmental variation is, therefore, important for understanding the stability of the interaction and the viability of the species that depend upon it (Heath and Tiffin, 2007; Thompson, 2005; Zoller et al., 2025). However, making such a description is not straightforward, especially in the case of highly specialized, interdependent interactions, in which one species may only be observed in close association with the other.

These problems come to a head in modeling the geographic distributions of interacting species. Most widely-used species distribution modeling (SDM) methods aim to predict the presence or absence of a species using environmental factors such as temperature, rainfall, and precipitation (Elith and Leathwick, 2009; Hill, 1991; Pereira and Itami, 1991); but they rarely incorporate the effects of other interacting species, much less model the environmental responses of predators, mutualists, or competitors (Lavergne et al., 2010; Treurnicht et al., 2016). Joint species distribution models (JSDMs) expand on traditional SDMs by modeling the presence of multiple species simultaneously, capturing shared environmental responses and potential biotic associations among them. Although this is an improvement, the biotic associations of JSDMs do not predict interactions, and the underlying mechanism of persistence driven by species interactions is still left unexplained (Poggiato et al., 2021).

A novel technique may let us address these issues in the specific case of specialized plant-pollinator mutualisms. In interactions where fruit production by a flowering plant is fully (or predominantly) dependent on the presence of animal pollinators, modeling flowering and fruit production in relation to annual variation in temperature and precipitation—rather than modeling a species’ presence in relation to multi-year averages—can identify conditions that support, first, flowering by the host plant and, second, the activity of its pollinators. This can be accomplished using data that is richer, but not dramatically much more effort-intensive, than presence records (Dakduk and Yoder, 2026; Yoder et al., 2024). The general approach of modeling annual reproductive activity rather than simple presence or absence, which we call temporal analysis of reproduction distributed in space (TARDIS), can capture relationships between climate and a species’s geographic distribution in much the same way as a presence-based SDM (Dakduk and Yoder, 2026), but it can also identify how population processes contribute to that relationship, and how they are affected by changing conditions (Yoder et al., 2024).

We first developed TARDIS modeling to understand how year-to-year weather variation influences masting by Joshua trees (*Yucca brevifolia* Engelm. and *Y. jaegeriana* McKelvey, Asparagaceae; Yoder et al. 2024). The two Joshua tree species are arborescent, succulent perennial monocots endemic to the Mojave Desert of southwestern North America. They are the largest plants in many Mojave plant communities, and are accordingly an important source of food and shelter for the region’s diverse fauna (DeFalco et al., 2010; Terrill et al., 2019; Yoder and Smith, 2024), as well as a cultural icon of the Mojave. Our TARDIS study of Joshua tree masting revealed that Joshua tree populations are more likely to mast when they experience a comparatively wet winter following a comparatively dry winter—and, because year-to-year variation in precipitation has increased since the early 20th century, the frequency of Joshua tree masting has also increased since that time (Yoder et al., 2024).

That study left a major question unanswered: how recent climate trends have impacted Joshua tree’s pollinators. Joshua trees are pollinated by highly specialized yucca moths (*Tegeticula antithetica* and *T. synthetica*, Prodoxidae). Yucca moths, genus *Tegeticula* (18 species) and *Parategeticula* (2) moths, are the exclusive pollinators of plants in the genera *Yucca* and *Hesperoyucca* (Davis, 1967; Pellmyr, 2003). Yucca moths use specialized mouthparts to actively pollinate the flowers of their host plants, after laying eggs in floral tissue, usually alongside developing ovaries. When the moth eggs hatch, the moth larvae then consume a portion of the seed crop. Yuccas require yucca moths for pollination, and do not self-pollinate; while host plants’ seeds are the moths’ only larval food resource (Pellmyr, 2003).

When yucca moths have completed their final larval instar, they emerge from the fruit in which they developed, drop to the ground, burrow into the soil to form a cocoon, and enter an extended diapause until the next flowering season. Little is known about the cues that trigger yucca moths to emerge from diapause, or even how long they can stay in diapause; it may be 29 years or longer, based on observations of *Prodoxus y-inversus*, a non-pollinating “bogus yucca moth” closely related to the Joshua tree pollinators (Powell, 2001). Like many desert perennials, Joshua trees do not flower every year—the TARDIS study estimate is that masting occurs once every four years (Yoder et al., 2024)—so it may be important that the moths judge their emergence precisely.

Many other insects use diapause to avoid unfavorable conditions, and the cues to emerge from diapause vary among species. Some species use changes in day length, temperature, rainfall, or even vibrations from nearby conspecifics (Denlinger, 2022; Hanski, 2026). Cold temperatures are often an indicator of given no other information, it might be expected that yucca moths emerge from diapause following the seasonal changes (Denlinger, 2022), allowing diapausing pupae to detect the passage of a year and emerge the next time conditions are right. Previous work on the yucca moth *Tegeticula yuccasella* and its host plant, *Yucca glauca*, indicated that moths emerge in two-year intervals that match the flowering frequency of their host plant (Fuller, 1990). Yet *T. yuccasella* also emerged when exposed to increased precipitation, even in off years from the two-year cycle (Fuller, 1990). The specific cues used by Joshua trees’ pollinators have not been directly studied, but we can reasonably expect they may not be identical to the factors that trigger host plant flowering.

Given no other information, it might be expected that yucca moths emerge from diapause following similar cues that prompt their host plants to flower. Our prior TARDIS model of flowering identified flowering as more likely in a year with more rainfall than the prior year, with warmer winter temperatures, and with relatively stable temperatures and drought stress over the prior two years (Yoder et al., 2024). The moths may respond to some or all of these weather cues, or they may deviate given the right stimulus (Fuller, 1990). They may emerge emerge without specific cues. If larvae simply vary in the duration of their diapause, it may be that larvae emerge into a good flowering year frequently enough to replenish the “moth bank” and carry the population through years in which moths emerge but Joshua trees do not flower.

Tracking yucca moths’ activity in any given year is challenging as they are small (~20mm), only active as adults during Joshua tree flowering, and do not consistently respond to light or scent traps (JBY, pers obs.); most recorded observations of Joshua tree’s pollinators have been taken in interaction with open flowers (Cole et al., 2017; Trelease, 1893). However, we can isolate yucca moths’ activity, to some degree, by tracking Joshua tree fruit set. Joshua trees only set fruit when the moths pollinate, so we can infer that if Joshua trees at a particular location produce fruit, yucca moths were present and active at that location. The converse is also true to some degree—and Joshua trees have been regularly observed to flower without producing fruit (PWM and JBY, pers obs; St. Clair and Hoines 2018; USA National Phenology Network 2026). Flowering without fruiting may reflect resource limitation as well as pollinator activity, but it is also true that a flowering tree cannot set fruit without pollinating yucca moths. We therefore hypothesized that, by modeling Joshua tree fruit production as distinct from flowering activity, we could isolate conditions that cue yucca moth emergence from diapause, to the extent they differ from conditions that cue Joshua tree flowering.

Here, we aim to better understand how weather variation may cue *Tegeticula antithetica* and *T. synthetica* emergence, and evaluate how recent climate trends in the Mojave may have impacted their populations. We accomplish this using a variant of the TARDIS analysis developed by Yoder *et al*. (2024), leveraging crowdsourced observations from the iNaturalist database to train separate models of annual flowering activity and fruiting activity in Joshua tree populations. We then compare the trained models to identify how conditions favoring fruiting differ from those favoring flowering, and hindcast predicted flowering and fruiting for each year from 1900 to 2025 to examine trends in flowering, fruiting, and potential mismatch between the two. Finally, we assess trends in predicted plant-pollinator mismatch in relation to elevation, latitude, and regions previously identified as climate refugia for Joshua tree, to examine how future climate change may impact the integrity of the Joshua tree-yucca moth mutualism.

## Methods

### Annotation and compilation of crowdsourced records

We obtained records of Joshua tree flowering and fruiting through the iNaturalist crowd-sourcing platform (inaturalist.org). Volunteer contributors to iNaturalist use a smartphone application or the platform’s website to upload photos of organisms with the time, date, and location of their observations (iNaturalist community, 2026). In our prior study modeling Joshua tree masting activity (Yoder and Smith, 2024), we confirmed species identity and annotated presence of flower buds, open flowers, fruits, or no evidence of flowering from images submitted with thousands of iNaturalist observations, using features of the iNaturalist online interface. We used functions from the package `rinat` to access the iNaturalist database through its API. We queried for “Research Grade” records, which have species identification confirmed by at least two iNaturalist contributors with at least 2/3 consensus on the identification, from the years 2008 (when iNaturalist launched) through 2025.

### Linking flowering records to weather data

We obtained monthly weather data from PRISM (PRISM Climate Group, Oregon State University, 2014), which provides spatially interpolated raster-formatted records of temperature (minimum and maximum, in degrees centigrade), vapor pressure deficit (minimum and maximum, in hPa) and total precipitation (in millimeters) across the contiguous U.S. from 1895 to the present. Following the process of Yoder *et al*. (2024), we aggregated iNaturalist records by year and grid cell within the ~4km raster grid of the PRISM data layers. Of the records within each grid cell in a given year, we determined whether any indicated flowering (annotated with flowers budding, open flowers, or fruit) and if so, we marked that grid cell as having *flowering present* in that year. If any records within the cell were annotated as having fruit, we additionally marked the cell as having *fruiting present* in that year. Finally, if all records within the cell were annotated as having no evidence of flowering, we marked it as having *flowering absent* and *fruiting absent*.

We matched each observation of annualized flowering and/or fruiting presence/absence with the corresponding weather records for up to two flowering years prior to the observation of flowering and/or fruiting. Joshua tree flowering typically begins in early spring and proceeds through late April, and we followed Yoder *et al*. (2024) in defining a flowering year as April through March of the following year; we considered possible weather predictors of flowering and fruiting from year 0 (the year of observation, Y0), back through year 1 (the year prior to observation, Y1), to year 2 (two years prior to observation, Y2). From PRISM data covering this time frame, we calculated quarterly total precipitation (mm), minimum and maximum temperatures (°C), and minimum and maximum vapor pressure deficit (hPa), designating quarters by the calendar: Q1, January–March; Q2, April–June; Q3, July–September; and Q4, October– December. We also calculated year-over-year differences in annual values (i.e., values for Y0 minus values for Y1 and values for Y1 minus values in Y2). The full list of candidate weather predictors is given in SI Table 1.

### Modeling flowering and fruiting activity

We trained Bayesian additive regression tree (BART) models predicting flowering activity in a given year and location (raster cell) with our candidate weather predictors. BART models are sum-of-tree classification and regression tree machine-learning models offering simplicity, flexibility, and comparative computational efficiency while at the same time providing ease of interpretation and transparent handling of model uncertainty (Carlson, 2020; Carlson et al., 2022; Chipman et al., 2010). We used functions in the `dbarts` (Dorie et al., 2014) and `embarcadero` (Carlson, 2020) packages to train and evaluate BART models predicting the presence or absence of flowering or fruiting in a given year, with the weather predictors described above. To model flowering activity, we effectively trained a model to differentiate weather associated with the presence of flower buds, flowers, or fruit from weather associated with observations of Joshua trees having no evidence of flowering; to model fruiting activity, we trained a model to differentiated weather associated with the presence of fruit from weather associated with the presence of flower buds or flowers.

We selected top predictors for each model from our longer list of candidate predictors using a procedure automated by `embarcadero`, which identifies the strongest predictors by training RI-BART models with 10, 20, 50, 100, or 200 component trees. Predictors with consistently higher frequency of inclusion in simpler models (i.e., models with fewer trees) are the most informative predictors, because simpler models are more strongly constrained to identify informative predictors (Chipman et al., 2010). We compared the final working models of flowering and fruiting activity in terms of the degree to which they shared top weather predictors — that is, whether the same factors contribute to flowering or fruit production — and the similarities of the posterior partial effects of those shared top predictors — whether those shared predictors shape the probability of flowering and fruiting in the same way. We then used our finalized models to hindcast flowering and fruiting for each year from 1900 to 2025, predicting reproductive activity using each model and the PRISM weather data for this period.

### Validation of modeled flowering and fruiting activity

To validate the models’ performance, we compared their predictions with independently collected records of flowering and fruiting activity and with observations from our own field surveys; these validation data comprised published reproductive surveys (St. Clair and Hoines 2018; n = 10), records from the National Phenology Network database (USA National Phenology Network 2026; USA-NPN, n = 195), annotations of flowers or fruits on digitized herbarium specimens (California Consortium of Herbaria 2024; Texas and Oklahoma Regional Consortium of Herbaria 2024; CCH, n = 197; TORCH, n = 27), digitized natural history records of moths (GBIF.org 2026; n = 20), and personal observation records by Christopher I. Smith (n = 136) and Ray Yeager (n = 19). With our field surveys (n = 172), this totaled 776 observations.

Observations of Joshua trees available from iNaturalist are distributed unevenly across the geographic range of the two species, concentrated in areas heavily trafficked by people (Fig. 2), and iNaturalist contributors rarely provide repeated observations of the same location over the course of the flowering season, that would let us identify whether specific populations that flower in a given year also produce fruit. To specifically test model predictions in geographic regions that are not well covered by iNaturalist contributors, and to provide observations of flowering with follow-up to check for fruit production, we conducted field surveys of Joshua tree flowering and fruiting in 2024 and 2025. Surveys consisted of two visits: the first early in the season (February—April) to record flowering presence/absence, and the second as a followup later in the season (April–June) to record fruit presence. At each survey site we walked 500 m transects, recording the number of trees that could be seen flowering or fruiting at 0, 250, and 500 m. We performed 60 spring-summer paired surveys in total, at 60 unique sites (Figure 2).

**Fig. 1.**
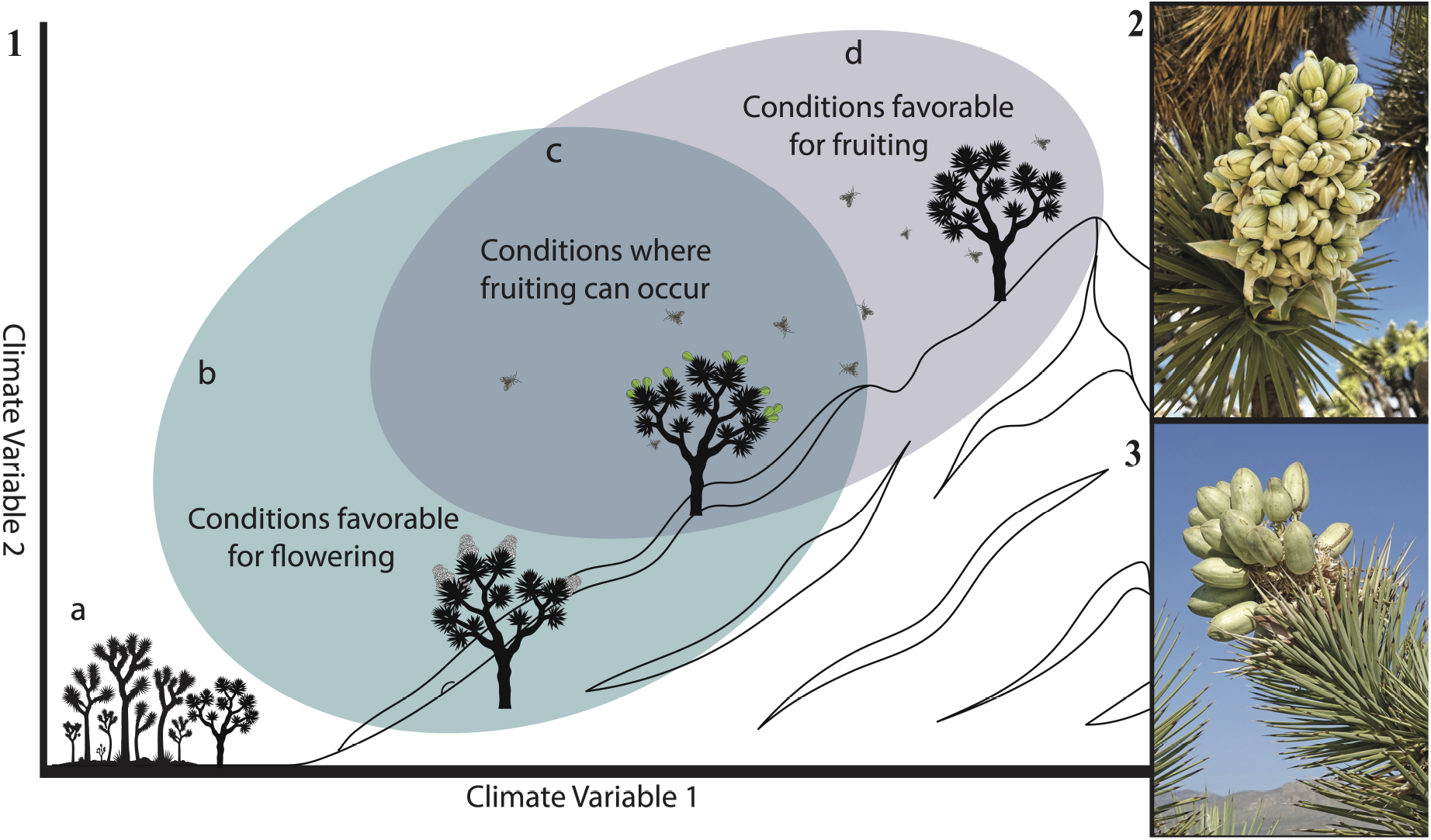
Conceptual diagram showing how climatic conditions structure Joshua tree reproductive activity and yucca moth emergence. A. No activity. B. Conditions favor flowering only. C. Conditions favor both fruiting and moth emergence. D. Conditions favor moth emergence only. 2. Western Joshua tree in flower photo by iNatualist user *jlboone2*. 3. Eastern Joshua tree bearing fruit photo by iNatualist user *sierra_the_seriema*.

**Fig. 2.**
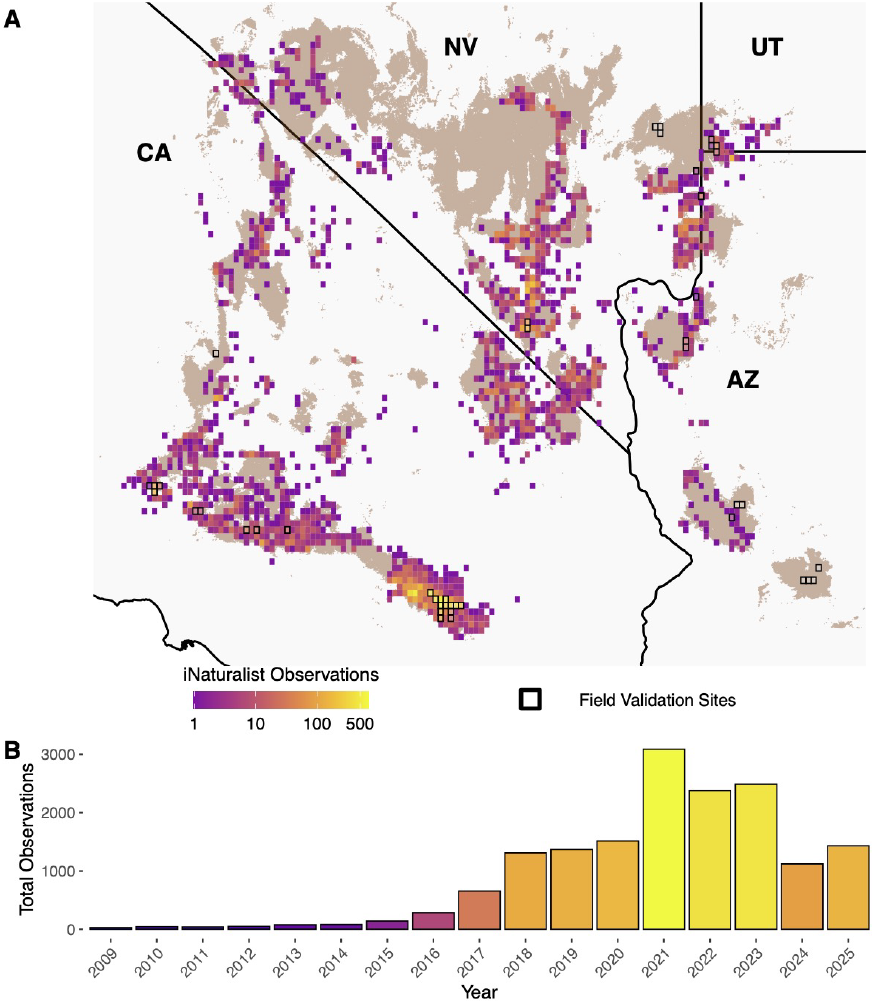
(A) The two Joshua tree species’ range (brown shading; Esque et al. 2023) overlaid with counts of total iNaturalist records (2008–2025) within cells of the PRISM raster layer grid. Locations surveyed in field in 2024-25 and used in model validation outlined in black. (B) Annotated Joshua tree iNaturalist records by year.

We assessed model accuracy by comparing the binary flowering or fruiting predictions of the models’ hindcast to the independent records we obtained from published papers and natural history records, and to our field survey observations, using a confusion matrix. This is a common type of contingency table used in machine learning that gives accuracy, precision, recall, and an F1 score, where accuracy reflects overall correct classifications, precision quantifies the correctness, and recall quantifies the completeness of positive predictions, and the F1 score provides a balanced measure of both. We additionally calculated AUC scores for each model on the validation data to be able to directly compare to the AUC of the training data. Finally, we used Welch’s t-tests to test whether our hindcast predicted higher probability of flowering or fruiting in years and locations where the validation records indicated flowering and fruiting were present, compared to years and locations where validation records indicated flowering and fruiting were absent.

### Testing for geographic and temporal trends in hindcast flowering and fruiting

To assess long-term trends in Joshua tree phenology, we generated annual predictions of flowering and fruiting probability across the species’ range for each year from 1900–2025 using our fitted BART models. For each year, we assembled the corresponding derived climate predictors. To quantify the direction and strength of change over time at each pixel, we calculated the Spearman rank correlation between year and predicted probability across the full 126-year time series, generating a spatial raster of trend strength and associated p-values for both flowering and fruiting. We then compared the magnitude of these trends using a paired t-test on the pixel-wise ρ value, testing whether the rate of increase differed significantly between flowering and fruiting across the landscape.

We examined trends in flowering and fruiting activity over the 126-year time span covered by our hindcast, and in relation to latitude, elevation, and climate refugia identified for Joshua trees in a recent high-resolution species distribution modeling study (Shryock et al., 2025). We tested for range-wide trends in flowering and fruiting by, first, estimating Spearman’s rank correlation between year and predicted probability of flowering or fruiting, for each raster cell falling within the range of the two Joshua tree species, using a range polygon derived from work by Esque *et al*. (2023).

We also converted the hindcast’s predicted probabilities of flowering and fruiting into binary predictions (using classification cutoffs determined by maximizing the true skill statistic; Carlson 2020) and summarized the frequency of years with flowering or fruiting across all years 1900–2025, and in 30-year periods in the earliest (‘historical’; 1900– 1929) and most recent (‘modern’; 1996–2025) portions of the hindcast period. We compared these historical and modern periods in terms of changing frequency of predicted flowering or fruiting years, and in terms of frequency of years with mismatch: either flowering predicted but not fruiting (i.e., years in which conditions favored flowering but not moth activity) or fruiting predicted but not flowering (years in which conditions may have favored moth activity, but not flowering).

We tested whether changes in the predicted frequency of flowering, fruiting, or mismatches were correlated with regions that should be climate refugia, as determined both *a priori* and by explicit species distribution modeling. We tested whether modeled changes in the frequency of, fruiting, and mismatch were correlated with elevation and latitude (because all else equal, we expect climate refugia to be at higher elevations and more northerly latitudes); and we examined these trends specifically within regions identified by Shryock *et al* (2025) as Joshua tree climate refugia in the late 21st century (2081-2100) under three different future climate scenarios of varying severity (SSP2-4.5, SSP3-7, and SSP5-8.; IPPC 2021).

## Results

### Data for model training and validation

We obtained 16,476 total iNaturalist phenology-annotated observations suitable as the training data for TARDIS analysis. The entire data set consisted of 13,142 observations with no evidence of flowering, 2,106 observations of flowering or flower budding, and 1,228 observations of fruiting. After aggregating all observations to the raster grid of PRISM data, we had 3,954 year-location flowering observations, and 1,124 year-location fruiting observations. Observations covered 32.65% of the total grid squares in the Joshua tree range (Fig. 2). Our field surveys during the 2024 and 2025 flowering seasons covered large portions of this range. Surveys recorded 179 flowering trees and 204 fruiting trees across 44 grid squares. We observed yucca moths on many flowering trees, but as many Joshua tree flowers were out of reach (more than 3m off the ground), confirmation of moth presence on all trees was not feasible in a comparatively simple survey protocol.

### BART models reveal contrasting conditions favoring flowering versus fruiting

Predictor selection using the method implemented in `embarcadero` (Carlson, 2020; Chipman et al., 2010) identified overlapping but not identical sets of most informative weather predictors for Joshua tree flowering and fruiting. The most informative predictors of flowering were, in order, Y1–Y2 contrast in precipitation (Δ_*Y*1*−*2_*PPT*), Y0–Y1 contrast in minimum temperature (Δ_*Y*0*−*1_ *MinTemp*), Y0–Y1 contrast in precipitation (Δ_*Y*0*−*1_*PPT*), precipitation in Q3 of Y0 (*PPT*_*Y*0*Q*3_), minimum vapor pressure deficit across Y1 (*MinVPD*_*Y*1_), and minimum temperature in Q4 (October–December) of Y0 (*MinTemp*_*Y*1*Q*4_) (Fig. S1). The most informative predictors of fruiting were, in order, minimum vapor pressure deficit in Q3 (July–September) of Y0 (*MinVPD*_*Y*0*Q*3_), the Y0–Y1 contrast in maximum temperature (Δ_*Y*0*−Y*1_ *MaxTemp*), minimum vapor pressure deficit across all of Y1 (*MinVPD*_*Y*1_), precipitation in Q3 of Y0 (*PPT*_*Y*0*Q*3_), maximum vapor pressure deficit in Q4 (October–December) of Y0 (*MaxVPD*_*Y*0*Q*4_), maximum temperature in Q4 of Y0 (*MaxTemp*_*Y*0*Q*4_) (Fig. S1).

Final working models trained with these predictors had good classification accuracy and fit for their training data (AUC = 0.81, RMSE = 0.55 for the flowering model; AUC = 0.76, RMSE = 0.55 for the fruiting model). The flowering model effectively predicts higher probability of flowering when the year of flowering (Y0) is wetter than the prior year (Y1), and the year prior (Y1) is drier than two years prior (Y2) ; when the year prior to flowering (Y1) has low drought stress; when the year of flowering (Y0) is cooler than the prior year (Y1); when temperatures in the fourth quarter of the year of flowering (Y0Q4) are above 0°C; and when the third quarter of the year of flowering (Y0Q3) has precipitation below 50 mm (Fig. 3). The fruiting model predicts higher probability of fruiting when the year of fruiting (Y0) is cooler than the year prior (Y1); when the year prior to fruiting (Y1) has low drought stress; when the third quarter of the year of fruiting (Y0Q3) has precipitation below 50 mm but also low drought stress; and when the fourth quarter of the year of fruiting (Y0Q4) has lower maximum temperatures and drought stress (Fig. 3).

**Fig. 3.**
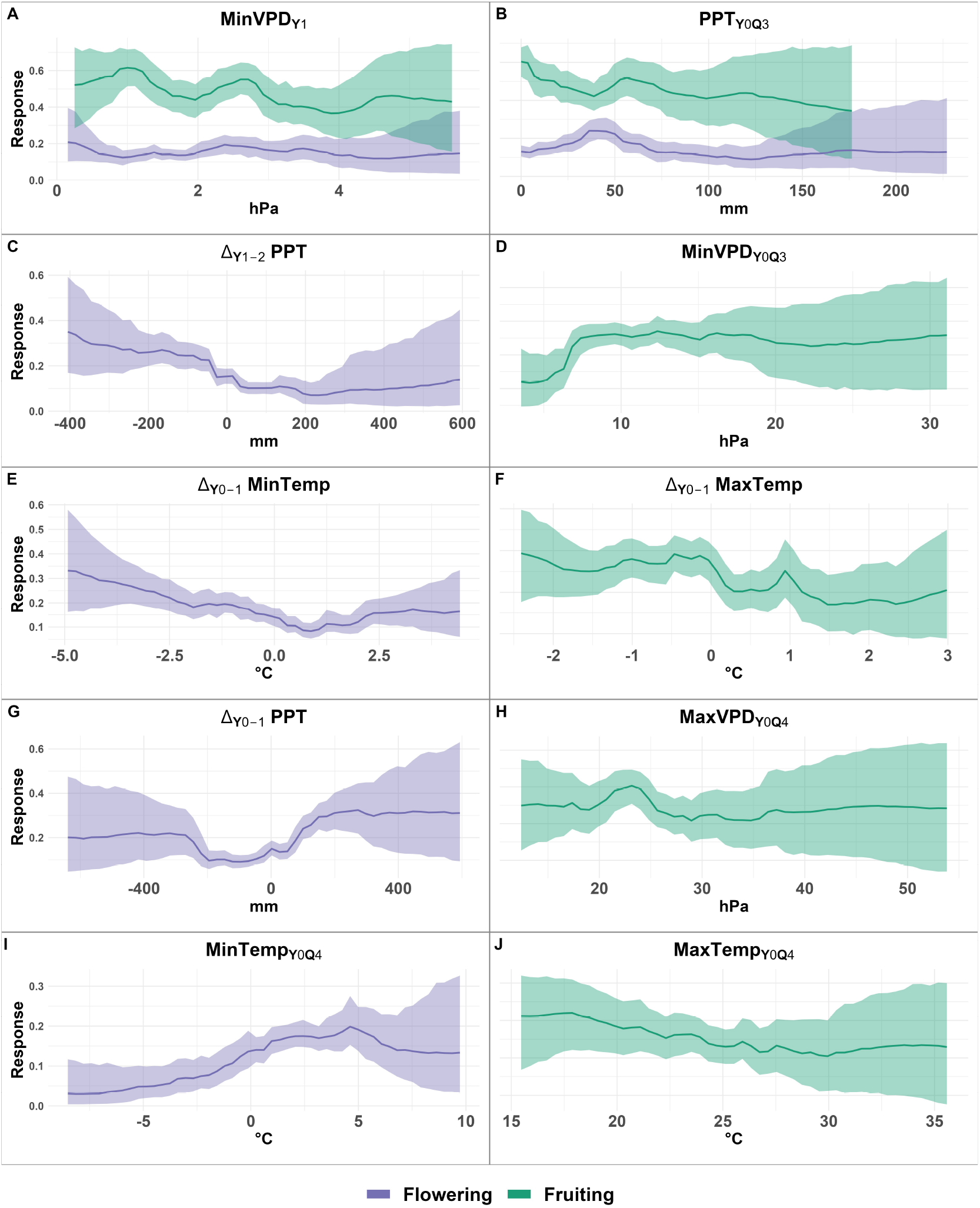
Posterior partial effects plots for the six most informative predictors of flowering (purple shaded areas and lines) and fruiting (green shaded areas and lines). In all panels, the x-axis is the value for a given predictor, and the y-axis is the marginal Pr(occurrence) for flowering or fruiting (or both); solid lines trace the median partial predictor effect, and shaded areas give the 95% posterior density distribution, from the final BART model.

Comparing the flowering model’s predictions to the field observations of flowering indicates moderate model performance (Accuracy = 0.49, Precision = 0.91, Recall = 0.39, F1 Score = 0.55); the model yields very few false positives but generates more false negatives. Similarly, comparing the fruiting model’s accuracy to the field observations of fruiting indicates moderate model performance (Accuracy = 0.52, Precision = 0.93, Recall = 0.4, F1 Score = 0.56); the model yields very few false positives but generates more false negatives. Validation against independent records and observations from our field surveys yielded AUC values of 0.687 (95% CI: 0.636 – 0.738) for flowering and 0.688 (95% CI: 0.63 – 745) for fruiting. Predicted flowering probability was higher at locations where flowering was observed in validation records (mean = 0.187) than where it was not (mean = 0.113; t-test, *t* = -6.73, *p <* 0.001). Predicted fruiting probability was also higher at locations where fruiting was observed (mean = .424) than where it was not (mean = 0.328; t-test, *t* = -6.21, *p <* 0.001).

### Range-wide trends in frequency of flowering and fruiting years

We predicted changes in the frequency of predicted Joshua tree flowering and fruiting over the 126 years our study period. The modeled probability of flowering increased over the time period (Spearman’s rank correlation test, modeled probability of flowering versus year: median *ρ* over all grid cells = 0.033, 95% CI 0.045 to 0.051); fruiting also increased slightly (median *ρ* = 0.0091, 95% CI 0.002 to 0.011). The rate of increase in flowering was significantly greater than the rate of increase in fruiting over the 126-year hindcast period (paired t-test, *t* = 13.94, *p <* 0.001, mean difference in *ρ* = 0.041, 95% CI 0.036 to 0.047).

We also compared predicted changes in the frequency of years with conditions favoring Joshua tree flowering and fruiting in 1996–2025 versus 1900–1929. Flowering predictions showed a higher frequency of flowering in the later period in 73.04% of grid cells. Across all cells in the species range, the flowering model predicted a median of two more years with flowering in 1996–2025 compared to 1900–1929 (mean = 2.18 more, 95% density of 3 fewer to 7 more). The fruiting model predicted a higher frequency of fruiting in the recent period in 42.07% of grid cells. The model predicted a median of zero more years with fruiting in 1996–2025 compared to 1900–1929 (mean = 0.439 fewer, 95% density of 9 fewer to 8 more; Fig. 4).

**Fig. 4.**
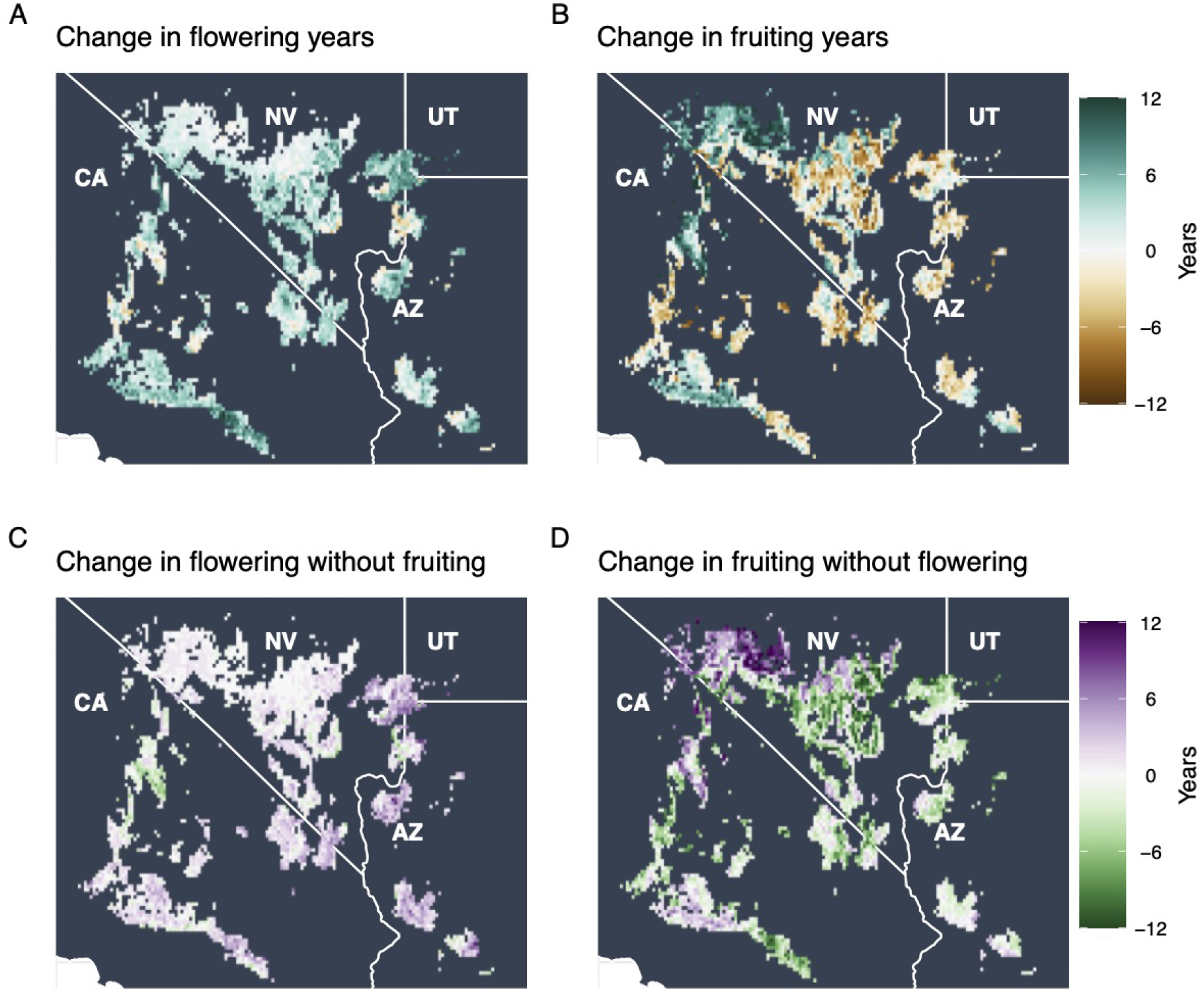
Predicted change in the frequency of years with conditions favoring flowering (flowering years) or fruiting (fruiting years), in the recent (1996-2025) versus early (1900-1929) 30-year periods. (A) Change in years with predicted flowering; (B) change in years with conditions suitable for flowering but not for fruiting; (C) change in years with conditions suitable for fruiting; (D) change in years with conditions suitable for fruiting but not for flowering.

We also examined changes in the frequency of predicted mismatch—years in which conditions were predicted to favor flowering but not fruiting, or fruiting but not flowering. We found the frequency of predicted flowering without fruiting increased across 52.17% of grid cells. Across the full range, the models predicted a median of one more year with flowering but no fruiting (mean = 0.94 more, 95% density of 4 fewer to 6 more). The frequency of years with predicted fruiting but no predicted flowering (i.e., years with conditions suitable for moth emergence and fruit production, but not for flowering) increased across 31.87% of grid cells. The models predicted a median of one less year with conditions favoring fruiting but no flowering (mean = 1.58 fewer, 95% density of 11 fewer to 7 more; Fig. 4).

### Changing frequency of flowering and fruiting years in climate refugia

Finally, we examined how these predicted changes in flowering and fruiting align with geographic regions that are either *a priori* climate change refugia (i.e., more northerly latitudes and higher elevations) or identified as refugia by formal species distribution models predicting suitable habitat for Joshua trees under future climate scenarios. We first tested whether change in years with conditions predicted to favor flowering, fruiting, or mismatches (Fig. 4) were correlated with latitude or elevation across the range of the two Joshua tree species. We found greater increases in the frequency of flowering years at higher elevation (Fig. 5A), but also at lower latitudes (Fig. 5B); there were significantly greater declines in the frequency of fruiting years at higher elevations (Fig. 5C) and at lower latitudes (Fig. 5D). These contrasting patterns meant that there was no significant correlation between change in the frequency of years suitable for flowering but not fruiting and either elevation or latitude (Fig. 5E,F); however there were significantly greater increases in the frequency of years suitable for fruiting but not flowering at lower elevations (Fig. 5G) and more northerly latitudes (Fig. 5H).

**Fig. 5.**
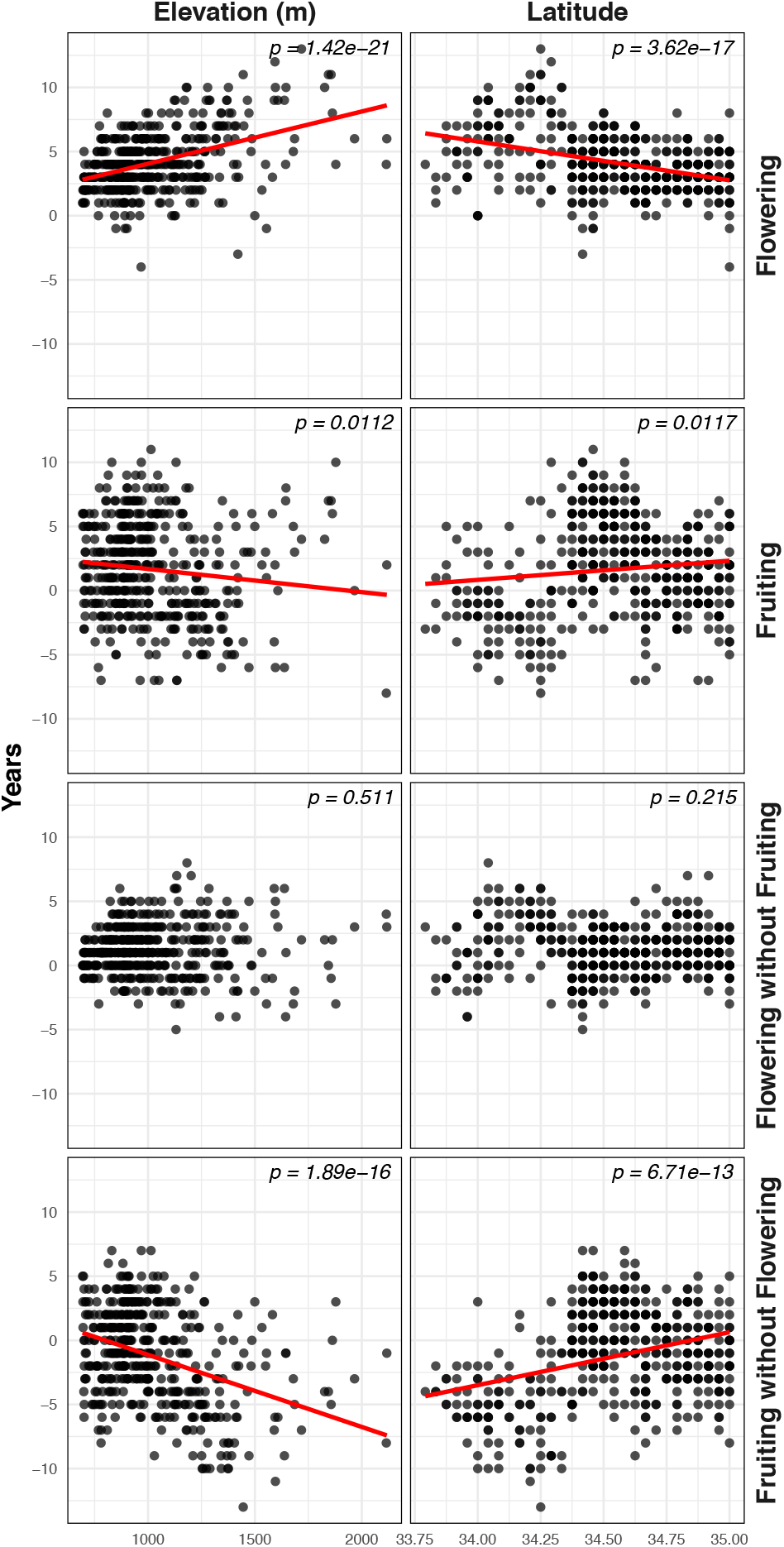
Predicted change in the frequency of years with conditions favoring flowering (flowering years) or fruiting (fruiting years), in the recent (1996-2025) versus early (1900-1929) 30-year periods, compared to *a priori* climate refugia.

We also compared the change in frequency of years with predicted flowering, fruiting, and mismatch conditions in areas that are predicted to be refugia for Joshua trees by 2100 under three different climate scenarios (Fig. S2; Shryock et al. 2025). The three scenarios represent low, medium, and high climate-forcing emissions; specifically SSP2– 4.5, SSP3–7, and SSP5–8.5, respectively, as defined by the 6th IPCC Assessment Report (Eyring et al., 2016). Trends in the frequency of flowering, fruiting, and mismatch years in areas identified as refugia under all three climate scenarios did not significantly differ from trends across the full species range (Fig. 6).

**Fig. 6.**
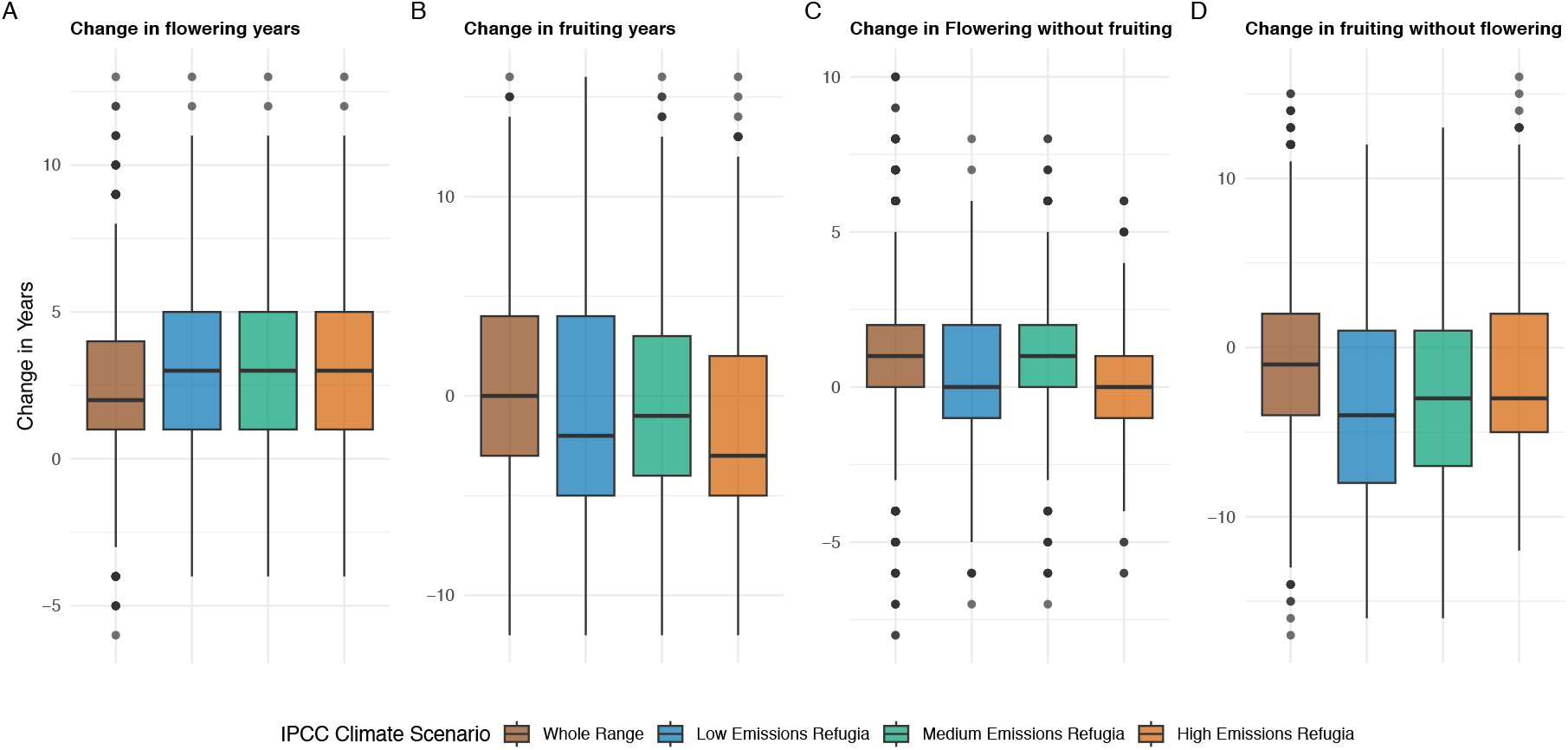
Change in frequency of years with conditions favoring flowering, fruiting, flowering but not fruiting, or fruiting but not flowering, from 1900–1929 to 1996–2025, across the full range of the two Joshua tree species (brown boxplots) and in regions identified as climate refugia for (2081–2100) under different three future climate scenarios (Fig. S2; blue, SSP2–4.5, green, SSP3–7, orange, SSP5–8.5). Box-and-whisker plots provide median (central bar), 25th and 75th percentiles (lower and upper box edges) and 1.5 *×* the interquartile range (whiskers) for change in years with predicted conditions across the specified areas.

## Discussion

The outcomes and stability of even closely coadapted mutualisms are often dependent on the broader environment in which mutualists interact (Heath and Tiffin, 2007; Pringle et al., 2013; Thompson, 2005). Disentangling how coadapted mutualists respond to environmental variation in time and space is challenging. Here, however, the specialized nature of the obligate pollination mutualism between Joshua tree and yucca moths has allowed us to begin to identify divergent responses of these mutualists to inter-annual weather variation, and to see how those differential responses may have impacted the effectiveness of the mutualism over more than a century of recent climate change.

By modeling flowering and fruiting separately, and comparing the predictor effects and predictions of the two models, we identify weather conditions that may cue pollinating yucca moths to emerge from diapause, and how they may differ from the weather triggers of Joshua tree masting. Hindcasting with our trained models of flowering and fruiting further reveals trends in conditions favoring these two elements of the yucca-moth mutualism over the past 125 years, a period during which average global temperatures have increased by more than 1°C (Calvin et al., 2023). This modeling helps us better understand how changing global climate may have altered the synchronicity of responses to annual variation in temperature and rainfall by Joshua trees and their pollinators— and thereby the resilience of this interdependent mutualism.

Modeling Joshua tree fruiting separately from flowering gives us an indirect view of the factors that cue *Tegeticula* yucca moths to eclose from diapause. It is possible that our models could be detecting the difference between the weather triggers of flowering and cues or resources needed for fruit formation. However, yucca moth pollination is necessary for fruiting (Fuller, 1990; Pellmyr, 2003; Trelease, 1893); though lack of fruiting may not mean yucca moths are absent, observations of fruit are proof positive of the moths’ presence and activity. The uncertainty inherent in using the presence of fruit as a proxy for moth activity is not unlike that in species distribution models based on records of presence or absence (Elith and Leathwick, 2009; Phillips et al., 2009): even directly observed absences reflect failures of detection as well as actual absences. With that caveat, we can cautiously interpret the differences between our models of flowering and fruiting as reflecting differences between weather cues for Joshua tree flowering and weather cues for yucca moth eclosion. The differences in the models’ predictions of trends over time should also reflect, to a similar degree, the differential responses of Joshua trees and their pollinators to 125 years of changing climate.

Our models also substantially simplify flowering and fruiting, treating each as a binary process—present or absent across an entire flowering year. This means we do not directly represent variation in the intensity of flowering or fruit production (or moth emergence), either between or within flowering years. It can be possible that many moths emerge, but only a few trees flower, which could reduce the total moth population if there are not enough flowers in which to rear larvae. Or, conversely, an intense flowering year may not be matched by moth emergence, so many trees flower without setting fruit even in conditions that support moth eclosion. Anecdotally, the intensity of moth and flowering activity in a given flowering year is often not well aligned at the local level (Christopher I. Smith, personal communication). Our models are also unable to address synchrony of Joshua tree flowering or moth eclosion within years; changes in the phenological synchrony of plants and key animal pollinators is a widely documented result of global climate change (Pausas, 2025; Piao et al., 2019; Zoller et al., 2025), but beyond the scope of what we can address here.

### Weather cues of flowering and fruit production

Our model of Joshua tree flowering recapitulates the results of Yoder *et al*. 2024, that Joshua tree flowering responds to contrasts in precipitation over multiple years, when a wetter year follows a drier year, and the year of flowering is cooler and has moderate to lower drought stress than the year before (Fig. 3). Many species of plants respond to contrasts in temperatures to trigger masting, not because it is a physiological necessity, but because it is a way to synchronize the mast with their fellow plants (Bogdziewicz et al., 2025; Kelly et al., 2013).

In our model of fruiting as a separate process from flowering, we find some of the same weather factors are top predictors; but these are two of the six most informative weather predictors in each model, and the shared predictors— minimum VPD in the year prior to flowering/fruiting and total precipitation in Q3 of the year of flowering/fruiting—have qualitatively different posterior partial effects (Fig. 3A,B). Beyond those two shared predictors, fruiting is associated with lower minimum and maximum VPD, and with cooler winter temperatures (*MaxTemp*_*Y*0*Q*4_; Fig. 3F) in a year that is cooler than the prior year (Δ_*Y*0*−*1_ *MaxTemp*; Fig. 3D); where flowering is associated with warmer winter temperatures (*MinTemp*_*Y*_0*Q*4; Fig. 3J).

Thus our BART modeling finds Joshua tree flowering responds to multi-year contrasts in precipitation, and to a lesser extent contrasts in temperature, while fruiting is more dependent on conditions in the months leading up to the flowering season, especially lower temperatures and drought stress (Fig. 3). (Though predictors of fruiting include change in annual maximum temperature; Fig. 3D.) For many diapausing insects, winter low temperatures are broadly important in cuing eclosion (Denlinger, 2022), so this is consistent with the possibility that our model of fruiting captures factors that trigger yucca moth eclosion.

Are the weather factors in our fruiting model the only cues yucca moths use to determine emergence from eclosion? Yucca moths might plausibly respond to a scent or chemical cue directly from their host trees. Joshua tree flowers do produce a unique scent profile that adult moths may use to locate flowers (Svensson et al., 2016). However, if floral scent were a primary eclosion cue for the moths, we would expect weather predictors of fruiting and flowering to be much more similar, if not identical. In the only direct study of yucca moth eclosion, *Tegeticula yuccasella* moths were observed to emerge from diapause underneath *Yucca glauca* plants that had not flowered, consistent with the hypothesis that yucca moths do not require cues from the closest host plants (Fuller, 1990). Moreover, other diapausing insects tend to emerge in response to weather conditions or day length (Denlinger, 2022; Poitou et al., 2020; Tachiki and Iwasa, 2013; Toxopeus et al., 2024). Direct experimental work could test whether the weather predictors we identified cause moths to emerge under controlled lab conditions, but given that the moths diapause for multiple years, up to decades, at a time, and may vary in their sensitivity to eclosion cues, it is challenging to design such experiments (Fuller, 1990).

### Projected increase in plant-pollinator mismatch with changing climate

Our models predict that the frequency of conditions favoring flowering has slightly increased since the early 20th century, while the frequency of conditions favoring fruiting has been stable to slightly decreasing. That is, Joshua trees are flowering somewhat more often without producing fruit, possibly because pollinating yucca moths are not emerging in concert with the increased frequency of flowering (Fig 4). This arises directly from recent trends in the weather predictors of flowering and fruiting. Climate change has likely already increased weather variability in the Mojave Desert, especially variability in annual precipitation: over the last decade, the region experienced the swung between the most intense drought in 1200 years and extreme precipitation events (Smith et al., 2023; Wahl et al., 2026; Williams et al., 2022, 2020). With Joshua trees responding to contrasts in precipitation, this could be leading them to flower more often. Meanwhile Joshua tree fruiting is associated with lower temperatures immediately before the flowering season (Fig. 3F,J), conditions that are harder to obtain as average and winter low temperatures rise.

These trends vary across elevational and latitudinal gradients, but not necessarily in ways that might be expected in *a priori* climate refugia at higher elevations and more northerly latitudes. As in the prior work by Yoder *et al*. (2024), we find predicted increases in the frequency of Joshua tree flowering are greater at higher elevations and at lower latitudes; but we see the opposite trends in predicted changes in the frequency of fruiting (Fig. 5). Predicted changes in the frequency of flowering without fruit production are not significantly correlated with elevation or latitude; but changes in the frequency of conditions favoring fruiting but not flowering are significantly negatively correlated with elevation and positively correlated with latitude (Fig. 5). That is, years in which our models predict moth emergence without Joshua tree flowering are becoming less common at higher elevations and in the southern part of the species’ range. These results contradict prior suggestions, based on admittedly limited observations, that cooler, higher-elevation habitats might be climate refugia for Joshua trees but not for their pollinators (Harrower and Gilbert, 2021).

We also examined modeled trends in conditions favoring flowering, fruiting, and both types of mismatch within regions identified as climate refugia in a recent species distribution modeling study based on exceptionally high resolution distribution data (Esque et al., 2023; Shryock et al., 2025). Modeled trends did not, however, significantly differ in regions identified as refugia in the late 21st century under low, medium, and high emissions scenarios, however; nor did trends in these regions differ from trends across the whole geographic range of the two Joshua tree species. That is, future climate refugia that we might identify *a priori* at higher elevations and more northerly latitudes; or those identified through formal distribution modeling by Shryock *et al*. (2025) have already experienced changes in flowering and fruiting conditions similar to the rest of the range.

### The Joshua tree pollination mutualism under changing climate

What does it mean for the viability of Joshua tree and yucca moth populations, and the stability of their mutualism, if they respond to changing climates differently, to a degree that mismatch—flowering without fruiting, or moth emergence without flowering—is increasing in frequency? Our results suggest a lower risk of the latter, but it is an open question how many consecutive years of moth emergence with a failure of flowering would be required to completely deplete the “moth bank” of diapausing larvae in the soil around Joshua tree populations. Some populations of *Tegeticula baccatella* moths that pollinate *Yucca baccata* have experienced collapse, and their host plants have failed to set fruit as a result. This was attributed to persistent herbivory of flowers by grazing livestock, which cut off the moths’ food source, though the duration of grazing needed to deplete moth populations was not determined (Lybbert and St. Clair, 2017). Yet, if persistent mismatches cause the localized extinction of moth populations, moths from the surrounding area may be able to recolonize. Anecdotal reports of cultivated Joshua trees setting fruit far from natural populations seem to indicate that yucca moths can disperse long distances and establish new populations (Christopher I. Smith, personal communication). This could allow the moths to recolonize after localized extinctions.

From the Joshua tree’s perspective, flowering without moths emerging would lead to failure of fruit set and seeds. More frequent occurrences of asynchrony are due to the trees flowering more often, without moths emerging more often. While Joshua trees are long lived and have many opportunities to reproduce over their lifespan, their seeds are short lived, and the conditions needed for seedling germination and survival are rare (Reynolds et al., 2012). Joshua tree seeds need a spring and summer with increased soil moisture, accompanied by warm soil temperatures, to germinate; and because they rapidly lose viability in the soil, they need these conditions soon after entering the seed bank. Given this, even minor disruptions to fruiting may jeopardize the few recruitment opportunities for Joshua trees may experience during their lifespan.

Other long-lived tree species in the American West face similar threats from climate change, in which mature forests persist as recruitment becomes less viable. In the Sierra Nevada mountains, 19.5% of conifer forests at low elevations are currently out of equilibrium with the climate (Hill et al., 2023). Models of ponderosa pine and Douglas-fir forests suggest that climate change will reduce recruitment at low elevations post-fire, and the forest edge will continue to move upslope (Kemp et al., 2019). Adult Joshua trees in areas with failing recruitment will continue to persist as long as there are no significant disturbances. Such Joshua tree populations may still support pollinator populations, and produce viable seed, even if little of it germinates, for some time to come. Further work to understand the population dynamics of yucca moths, how long they can stay in diapause, and their typical dispersal distances can more directly assess the risks mismatches may pose.

### Conclusions

Our study is the first to present evidence of the weather cues that yucca moths use to trigger emergence from eclosion to pollinate Joshua trees. Our models predict an increasing frequency of mismatch between flowering trees and moth emergence, in which conditions favor Joshua tree flowering but not fruit production that depends on yucca moth emergence. While yucca moths may be able to recolonize areas if repeated mismatches cause local population collapses, potential pollen limitation could reduce the already scant recruitment opportunities for Joshua trees, including in regions that may otherwise be future climate refugia. These findings underscore the vulnerabilities of tightly coevolved interactions to environmental change, and highlight the need for further research on yucca moth diapause dynamics and dispersal to inform conservation strategies for these Mojave Desert icons.

## Acknowledgments

We thank the iNaturalist contributor community and the USA National Phenology Network and contributors to its *Nature’s Notebook* program, for making this work possible. Support was provided by CSU Northridge (to PWM and JBY), by the Desert Symposium Reynolds Award, Judith Presch Research Scholarship, and Marian Huff Tremblay Science Scholarship at Joshua Tree National Park (to PWM); and by the U.S. Bureau of Land Management (to JBY, L24AC00509-00), and by the U.S. National Science Foundation (to JBY, DEB 2001180 and 2536007). Evan Jabbaz, Daniel Dakduk, Robert Avila assisted with fieldwork and data annotation, and Colin J. Carlson and Christopher I. Smith provided helpful comments on analysis design and interpretation.

## Data accessibility

Code for the acquisition and management of training data, and for running all analyses, is available for review on GitHub: github.com/skymasterpryce-droid/Joshua-Tree-Fruiting-Moth-Code

## Author contributions

Both authors contributed to conceiving and designing the project; PWM curated iNaturalist records, managed model development, and conducted fieldwork for model validation; PWM drafted the manuscript and both authors contributed to editing.

## Supporting information

**Fig. S1.**
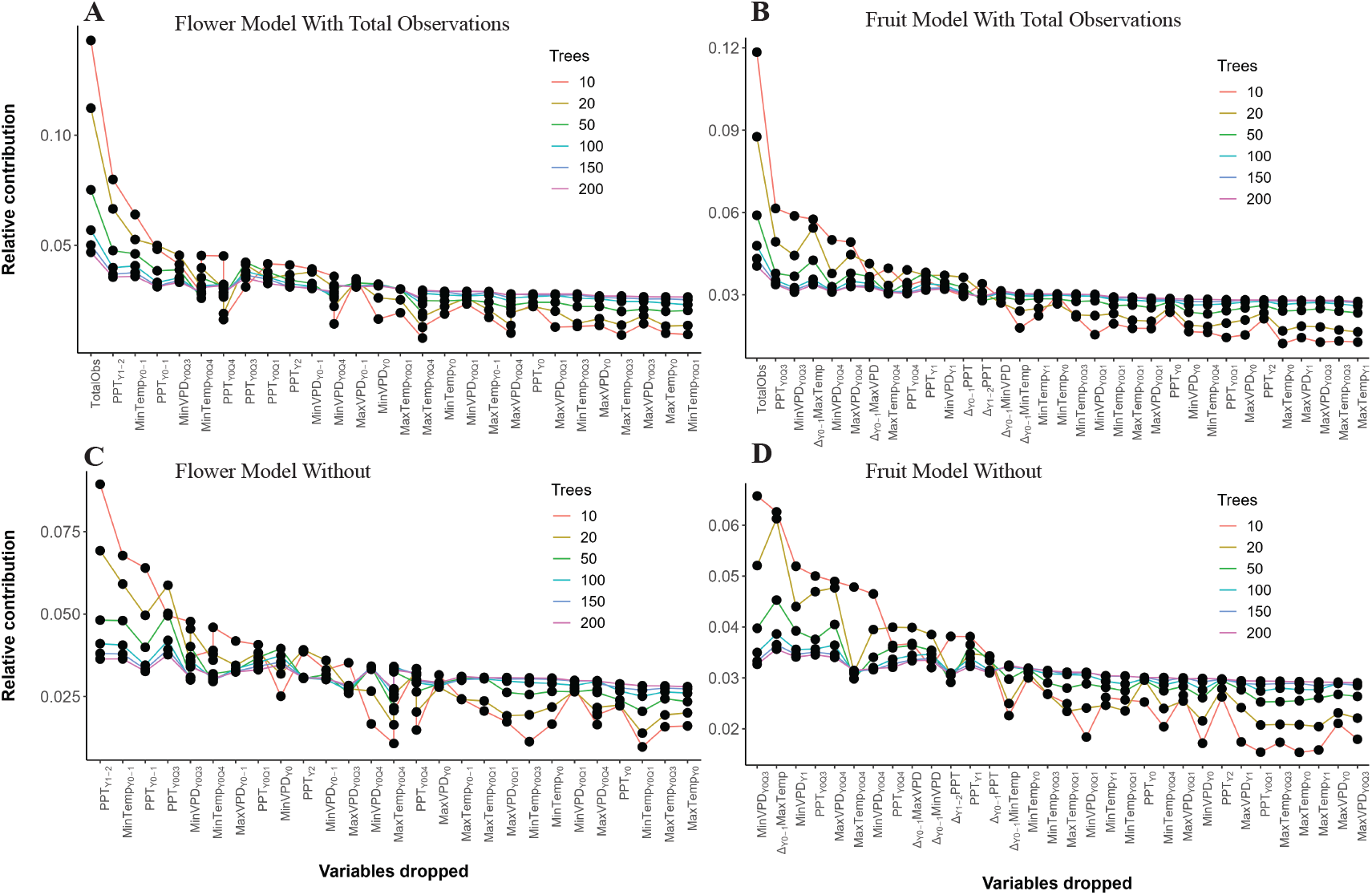
Predictor selection for BART models of Joshua tree flowering and fruiting.

**Fig. S2.**
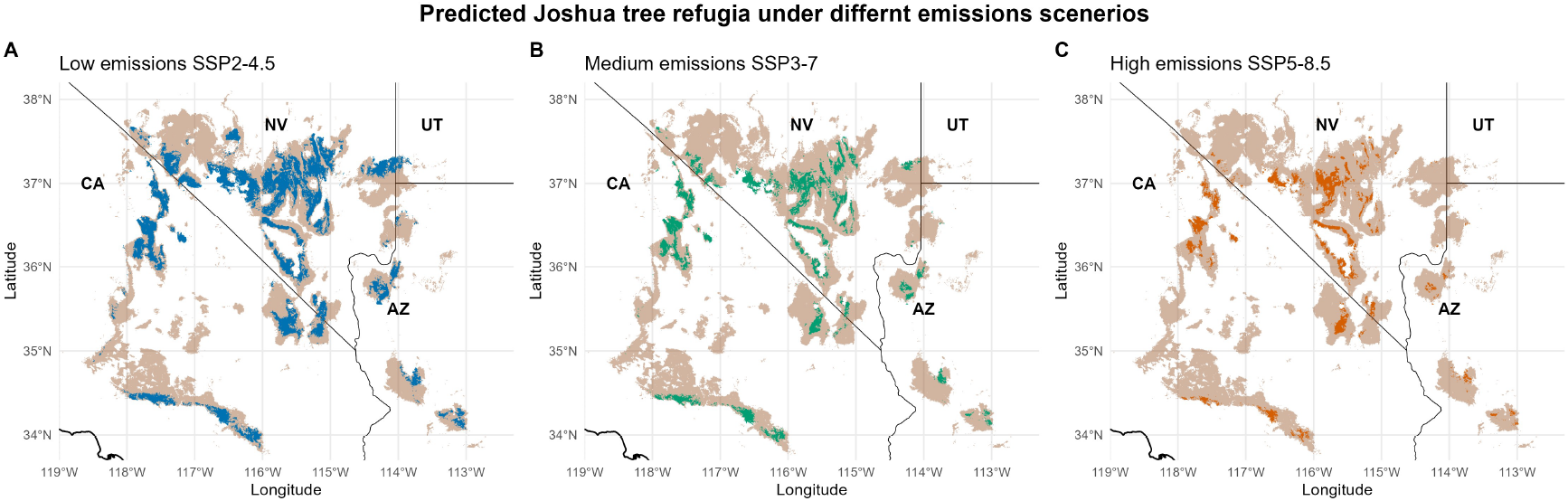
Spatial extent of predicted climate refugia for Joshua tree in 2081-2100 (??) under three emissions scenarios, as modeled by Shryock *et al*. (2025).

